# A modular plasmid toolkit applied in marine Proteobacteria reveals functional insights during bacteria-stimulated metamorphosis

**DOI:** 10.1101/2023.01.31.526474

**Authors:** Amanda T. Alker, Alpher E. Aspiras, Tiffany L. Dunbar, Morgan V. Farrell, Andriy Fedoriouk, Jeffrey E. Jones, Sama R. Mikhail, Gabriella Y. Salcedo, Bradley S. Moore, Nicholas J. Shikuma

## Abstract

A conspicuous roadblock to studying marine bacteria for fundamental research and biotechnology is a lack of modular synthetic biology tools for their genetic manipulation. Here, we applied, and generated new parts for, a modular plasmid toolkit to study marine bacteria in the context of symbioses and host-microbe interactions. To demonstrate the utility of this plasmid system, we genetically manipulated the marine bacterium *Pseudoalteromonas luteoviolacea*, which stimulates the metamorphosis of the model tubeworm, *Hydroides elegans*. Using these tools, we quantified constitutive and native promoter expression, developed reporter strains that enable the imaging of host-bacteria interactions, and used CRISPR interference (CRISPRi) to knock down a secondary metabolite and a host-associated gene. We demonstrate the broader utility of this modular system for rapidly creating and iteratively testing genetic tractability by modifying marine bacteria that are known to be associated with diverse host-microbe symbioses. These efforts enabled the successful transformation of twelve marine strains across two Proteobacteria classes, four orders and ten genera. Altogether, the present study demonstrates how synthetic biology strategies enable the investigation of marine microbes and marine host-microbe symbioses with broader implications for environmental restoration and biotechnology.

## INTRODUCTION

Marine bacteria are a valuable and currently under-utilized resource for environmental restoration (1–6) and bioprospecting (7, 8), especially considering their influence on biogeochemical cycles (9) and their vital role in evolution through symbioses with eukaryotes (10). While advances in metagenomic sequencing have enabled a deep exploration of microbial diversity and gene content (11, 12), genetic tools to explore functions in marine bacteria remain scarce.

Effective genetic engineering approaches in model microbial species, such as *E. coli*, utilize standardized and modular cloning toolkits (13–19), which leverage aligned plasmid parts based on the ordered pairings of restriction site overhangs to enable innumerable mix-and-match plasmid assembly options. However, such modular genetic tools have not yet been applied to most marine bacterial species. Thus, adapting and applying standardized molecular cloning tools for studying marine bacteria can provide a framework for addressing functional questions for basic science and biotechnology.

Marine Proteobacteria are of specific interest as targets for genetic tool development due to their ability to produce diverse bioactive metabolites (20), their prominent associations in aquatic microbiomes, and involvement in host-microbe symbioses (21–23). Alphaproteobacteria and Gammaproteobacteria, in particular, are the most abundant orders in the ocean (12) and are prominent members of the microbiomes of animals such as phytoplankton (12), tubeworms (21) and corals (24). However, the vast majority of environmental strains have not been interrogated using a genetics approach, leaving our ability to manipulate marine microbes limited to a few representative strains.

Of particular interest as targets for genetic manipulation are marine *Pseudoalteromonas* species because they produce a number of bioactive secondary metabolites (8, 25–29) and are often found in association with marine invertebrates (30–36). *Pseudoalteromonas* species are known to engage in a transient symbiosis called bacteria-stimulated metamorphosis, whereby surface-bound bacteria promote the larval-to-juvenile life cycle transition in invertebrates such as tubeworms and corals (37, 38). *Pseudoalteromonas luteoviolacea* stimulates the metamorphosis of the tubeworm *Hydroides elegans* (39, 40) by producing syringe-like protein complexes called Metamorphosis-Associated Contractile structures (MACs). MACs stimulate tubeworm metamorphosis by injecting an effector protein termed Mif1 into tubeworm larvae (40–42). Genes encoding the MACs structure are found in the *P. luteoviolacea* genome as a gene cluster encoding structural components, such as the *macB* baseplate and *macS* sheath, as well as the protein effector gene *mif1* (41). Despite the significant insights gained by using genetics in *P. luteoviolacea*, new genetic tools are needed to further dissect the function of MACs and their stimulation of tubeworm metamorphosis.

In this work, we utilize a modular plasmid toolkit, and contribute new Marine Modification Kit (MMK) plasmid parts, to study bacteria-stimulated metamorphosis in the Gammaproteobacterium, *P. luteoviolacea*. We demonstrate the broader utility of this plasmid system by manipulating marine Alphaproteobacteria and Gammaproteobacteria that have been shown previously to be involved in diverse host-microbe interactions.

## RESULTS

### Toolkit-enabled quantitative promoter expression in P. luteoviolacea

To test the application of modular genetic tools in marine bacteria, we identified a set of preexisting parts from the Yeast Toolkit and Bee Toolkit platforms (17, 18) and used Golden Gate Assembly (14) for rapid, modular construction of plasmids (Figure 1A-C). Each type of part is defined by its functional role (e.g. promoter, coding sequence) and directional 4 bp overhangs generated by flanking Type IIS (BsaI) restriction sites. The modular parts include: Type-1 and Type-5 stage-2 connectors with BsmBI recognition sites (17, 18), a Type-2 promoter with ribosome binding site (RBS), a Type-3 protein coding sequence (CDS), a Type-4 terminator, an optional Type-6 repressor and Type-7 promoter with RBS, and a Type-8 backbone. For this work, we selected a broad-host-range (BHR) plasmid backbone containing a kanamycin resistance gene, a reporter coding sequence (fluorescent *gfp-*optim1, *mRuby* or *Nanoluciferase* [*Nluc*]), T7 terminator and a stage-2 assembly connector. The backbone has an RSF1010 origin of replication, known to replicate in a broad range of gram-positive and negative bacterial hosts (43). A promiscuous origin of transfer and plasmid-encoded conjugative machinery (44) enabled domestication-free conjugative transfer with MFD*pir* auxotrophic host *E. coli* cells (45).

**Figure 1.**
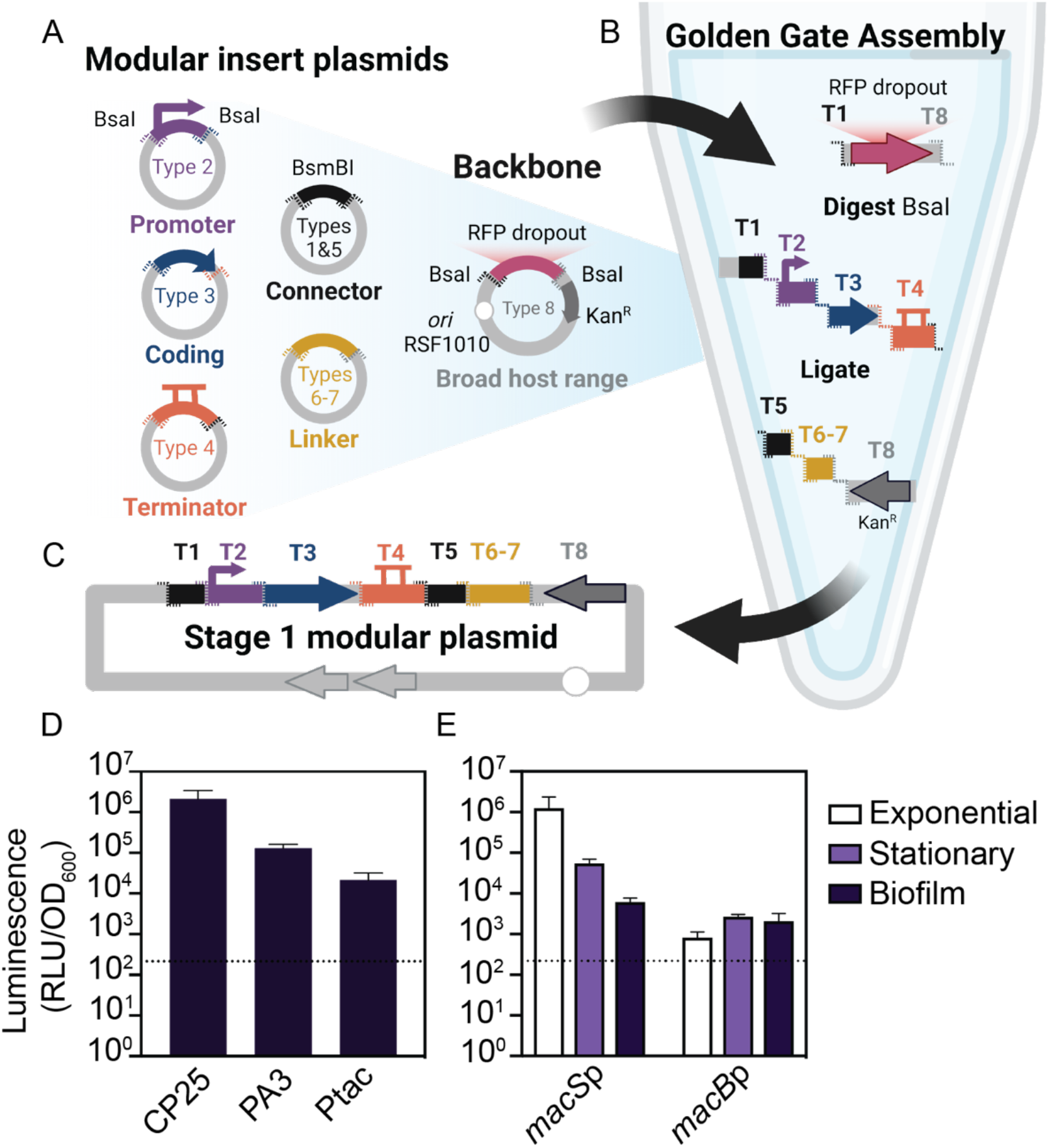
Schematic overview of the modular plasmid system and quantitative promoter measurements. (**A)** Schematic representation of the modular golden gate assembly plasmid parts with flanking BsaI cut sites (dashed lines). Overlapping 4 bp overhangs are color coordinated. The modular broad host range (BHR) backbone (pBTK402) contains inverted BsaI cut sites and an RFP dropout. **(B)** Golden Gate Assembly is performed in a one-tube reaction by digesting the backbone and insert part plasmids with BsaI and ligating with T4 ligase. **(C)** A modular stage-1 plasmid is complete when all overlapping inserts are successfully assembled in order. **(D)** Biofilm luciferase assay of *P. luteoviolacea* strains expressing plasmids with different constitutive promoters driving a Nanoluciferase (*Nluc*) gene (CP25-*Nluc*-T7, PA3-*Nluc*-T7, Ptac-*Nluc*-T7). Luminescence, as relative luminescence units (RLU), is normalized to optical density at 600 nm (OD_600_) and plotted on a log base 10 scale. The dashed line indicates the detection limit three standard deviations above the *P. luteoviolacea* (no plasmid) control (Y= 214 RLU/OD_600_). Plotted is the mean of three biological replicates. Error bars indicate standard deviations. **(E)** Luciferase assay comparing native MACs *macS* and *macB* promoters linked with a *Nluc* coding sequence across different modes of growth. N=3 biological replicates. Error bars indicate standard deviations. The dashed line indicates the detection limit three standard deviations above the *P. luteoviolacea* (no plasmid) control (Y= 218 RLU/OD_600_).

To apply the modular genetic tools in a marine symbiosis model, we explored constitutive and native promoter expression in *P. luteoviolacea*. We assembled plasmids with one of five promoters fused to *Nluc* and conjugated the plasmids into *P. luteoviolacea*. We utilized two existing constitutive promoters, PA3 and CP25, previously shown to work in diverse bee gut microbes (17, 46, 47). We designed a Ptac LacO constitutive promoter part (pMMK201), which is a hybrid of the *lac* and *trp* promoters amplified from the pANT4 plasmid (48). When *P. luteoviolacea* with the plasmids were grown as a biofilm, we observed at least 10-fold more luminescence signal compared to the background with all constitutive promoters tested (Figure 1D). The CP25 promoter exhibited a 10,000-fold increase in luminescence. We also constructed two native *P. luteoviolacea* promoters driving the expression of the MACs structural genes; promoters from the MACs sheath (*macS* promoter, pMMK203) and baseplate (*macB* promoter, pMMK202) genes. The *macS*p luciferase reporter strain was elevated 1,000-fold in exponential growth as compared to 100-fold in stationary and 10-fold in biofilm phase, when compared to the detection limit (Figure 1E). In contrast, the *macB*, baseplate promoter exhibited similar levels of luminescence among each phase, approximately 10-fold higher than the detection limit (Figure 1E).

### Functional CRISPRi knockdown of secondary metabolite biosynthesis in P. luteoviolacea

While previous studies in *P. luteoviolacea* have used gene knockouts to interrogate gene function, these approaches are time consuming and low-throughput. We therefore tested whether *P. luteoviolacea* is amenable to gene knockdown via CRISPR interference (CRISPRi) (Figure 2A and B) (49, 50). As a proof-of-concept, we targeted the *vioA* gene that encodes a key enzyme in the biosynthesis of violacein (51), which gives *P. luteoviolacea* its characteristic purple pigment (Figure 2B). To facilitate assembly for and expression in *P. luteoviolacea*, we modified the BsmBI cut site in the dCas9 part plasmid to include the *bla* gene (pMMK601), thus also conferring resistance to ampicillin. We replaced the existing PA1 promoter with Ptac in the guide RNA part plasmid targeting *gfp* (pMMK602). An assembled plasmid containing dCas9 and a single guide RNA (sgRNA) targeting the non-template strand of *vioA* (pMMK603) was conjugated into *P. luteoviolacea* resulting in the visible absence of the purple pigment associated with violacein production on the plate (Figure 2C). *P. luteoviolacea* with a sgRNA targeting *gfp* was included as a non-targeting control. A significant reduction of violacein production was observed between cultures of *P. luteoviolacea* strains expressing the *vioA* and *gfp* targeting CRISPRi plasmids (p=0.0007, Figure 2D). The lack of violacein in the *vioA* knockdown strain was comparable to that of a *P. luteoviolacea* strain with an in-frame deletion of *vioA* (Figure 2D). These results demonstrate the successful implementation of CRISPRi for gene knockdown in *P. luteoviolacea*.

**Figure 2.**
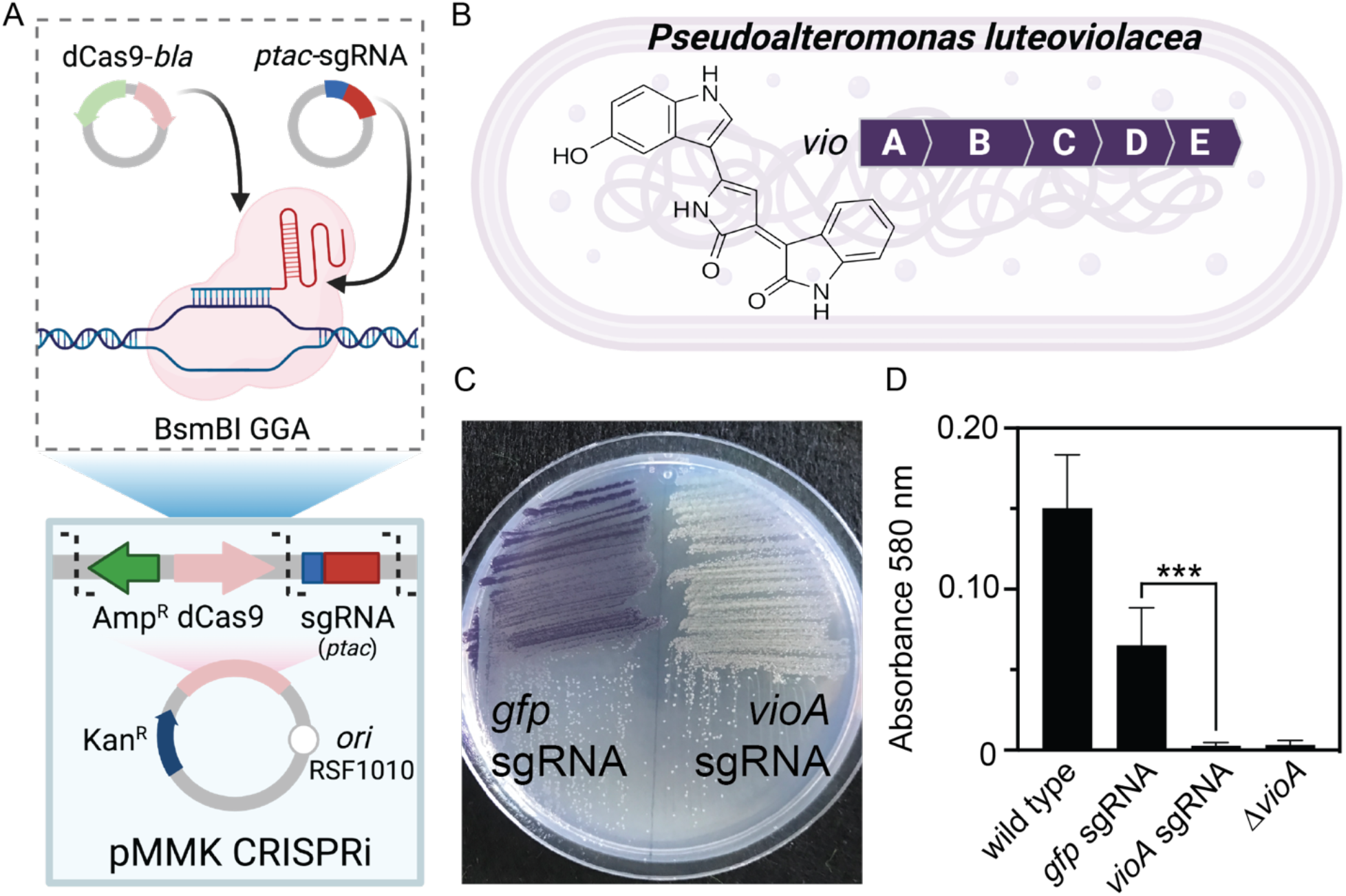
CRISPRi knockdown of secondary metabolite production in *P. luteoviolacea*. **(A)** Schematic representation of modular CRISPRi parts adapted to include dCas9-*bla* and Ptac-sgRNA parts, pMMK601 and pMMK602, respectively. Part plasmids are combined and a Golden Gate Assembly was performed with BsmBI. **(B)** Schematic representation of the violacein genecluster *vioABCD* in *P. luteoviolacea* and the violacein molecular structure. The CRISPRi system was assembled with an sgRNA targeting the *vioA* gene (pMMK603) and employed to knock down violacein production in *P. luteoviolacea*. **(C)** *P. luteoviolacea* with *gfp* (pMMK602) or *vioA* (pMMK603) sgRNA plasmids grown on marine agar plates. **(D)** Quantification of violacein production (measured at 580 nm) between *P. luteoviolacea* containing *gfp* or *vioA* sgRNA plasmids. Asterisks indicate significant differences (***p=0.0007, Dunnett’s T3 multiple comparisons test). *P. luteoviolacea* wild type and ∆*vioA* strains are included as controls. Bars represent the mean (N=8) and error bars indicate standard deviations.

### Functional CRISPRi knockdown and visualization of P. luteoviolacea during a tubeworm-microbe interaction

We next tested whether CRISPRi would be functional in the context of a marine host-microbe interaction by targeting the *macB* gene, which encodes the MACs baseplate, an essential component of the MACs complex that induces tubeworm metamorphosis (39, 40) (Figure 3A). Biofilm metamorphosis assays were performed comparing *P. luteoviolacea* strains with sgRNAs targeting *macB* (pMMK604) or a sgRNA targeting *gfp* as a control (Figure 3B). The strain containing the *macB* sgRNA exhibited significantly reduced levels of tubeworm metamorphosis compared to the *gfp-*sgRNA control (Figure 3B; Mann Whitney test, p=0.029). The reduction of metamorphosis stimulation in the *macB*-sgRNA knockdown strain was comparable to that of a *P. luteoviolacea* strain with an in-frame deletion of *macB* carrying the *gfp-*sgRNA control plasmid (Figure 3B). These results demonstrate that CRISPRi paired with a modular plasmid system is a viable tool for interrogating gene function during a marine host-microbe interaction.

**Figure 3.**
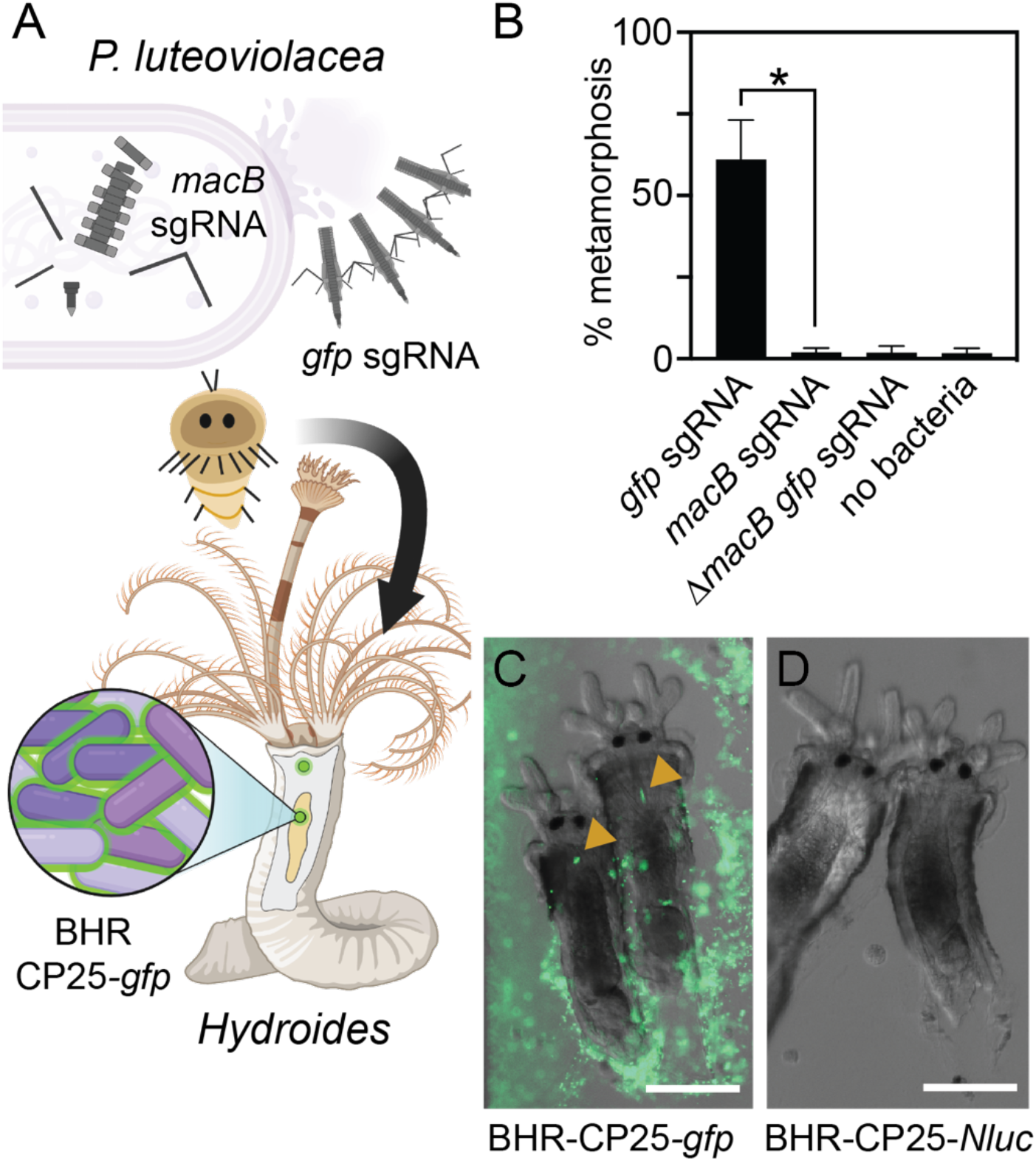
Functional knockdown of MACs and visualization of *P. luteoviolacea* during the tubeworm-microbe interaction. (A) Schematic depicting *P. luteoviolacea* and the production of MACs, which induce tubeworm metamorphosis. CRISPRi single guide RNA (sgRNA) targeting the *macB* MACs baseplate gene prevents MACs from assembling, rendering the bacterium unable to induce metamorphosis. Cells that produce intact MACs are able to induce tubeworm metamorphosis. A strong fluorescent reporter strain (BHR-CP25-*gfp*) enabled visualization of live tubeworm-bacteria interaction. (B) Bar graph representing biofilm metamorphosis assays with *P. luteoviolacea* carrying a CRISPRi plasmid targeting *macB* or *gfp* and *Hydroides* tubeworms. A *P. luteoviolacea* ∆*macB* strain with a sgRNA targeting *gfp* and a treatment without bacteria (no bacteria) were included as controls. Biofilm concentrations were made with cells at OD_600_ 0.2. Bars plotted are the average of 3 biological replicates (N=3) performed on separate occasions. Four technical replicates were performed for each treatment during each biological replicate, with each well containing 20-40 worms. Error bars indicate standard deviations. Statistical significance between treatments is indicated by an asterisk (*p=0.029, Mann Whitney test). (C and D) Fluorescence micrographs of *Hydroides elegans* juveniles imaged 24 hours after the competent larvae were exposed to inductive biofilms of *P. luteoviolacea* containing plasmids with (C) CP25-*gfp* or (D) CP25-*Nluc*. Strains containing *Nluc* plasmids were used as a negative control to account for autofluorescence. Yellow arrows show accumulation of fluorescent bacteria in the *Hydroides* juvenile pharynx. Scale bar is 100 µm.

To date, bacteria have not been visualized during or after the stimulation of metamorphosis in *Hydroides*. To test whether marine bacteria harboring a toolkit plasmid are amenable to live cell imaging when in association with juvenile tubeworms, we created biofilms of *P. luteoviolacea* containing plasmids encoding CP25-*gfp-*T7 (*gfp*) or CP25-*Nanoluc-*T7 (*Nluc*) and added competent *Hydroides* larvae. After incubation for 24 hours, biofilms of *gfp*-expressing *P. luteoviolacea* were clearly observed when visualized by fluorescence microscopy (Figure 3C). *P. luteoviolacea* stimulated *Hydroides* metamorphosis while carrying a modular plasmid and fluorescent bacteria were observed being ingested by the *Hydroides* juveniles. Bacteria can be seen collecting in the pharynx (Figure 3C, yellow arrows), then moving in a peristaltic fashion toward the gut (Movie S1). In contrast, bacteria and their biofilms were difficult to visualize by light microscopy without fluorescent bacteria (Figure 3D). Taken together, the modular plasmid system enables live imaging and experimentation during a marine host-microbe interaction.

### Genetic manipulation of diverse marine Proteobacteria

Given the success of genetic manipulation of *P. luteoviolacea*, we tested whether more diverse marine Proteobacteria are amenable to genetic manipulation via the modular genetic toolkit. To this end, we isolated or acquired representative bacteria that are known to engage in symbioses with marine plants or animals in the ocean (Figure 4A; Table S1). To enable genetic selection using antibiotics, we determined the minimum inhibitory concentration for each bacterial strain tested against kanamycin (Table S1). When conjugation was performed using the broad-host-range (RSF1010) plasmid backbone, CP25 promoter, *gfp* reporter and T7 terminator, we observed the expression of *gfp* in 12 marine strains across two proteobacterial classes, four orders and 10 genera (Figure 4B). Eight of the strains were made tractable for the first time, including bacteria from *Pseudoalteromonas, Endozoicomonas, Cobetia, Shimia, Nereida, Leisingera*, and *Phaeobacter* genera (Figure 4B). Adaptations to the conjugation protocol and use of constitutive promoters driving *gfp* enabled visual confirmation of successful conjugation (Figure 4B, Methods).

**Figure 4.**
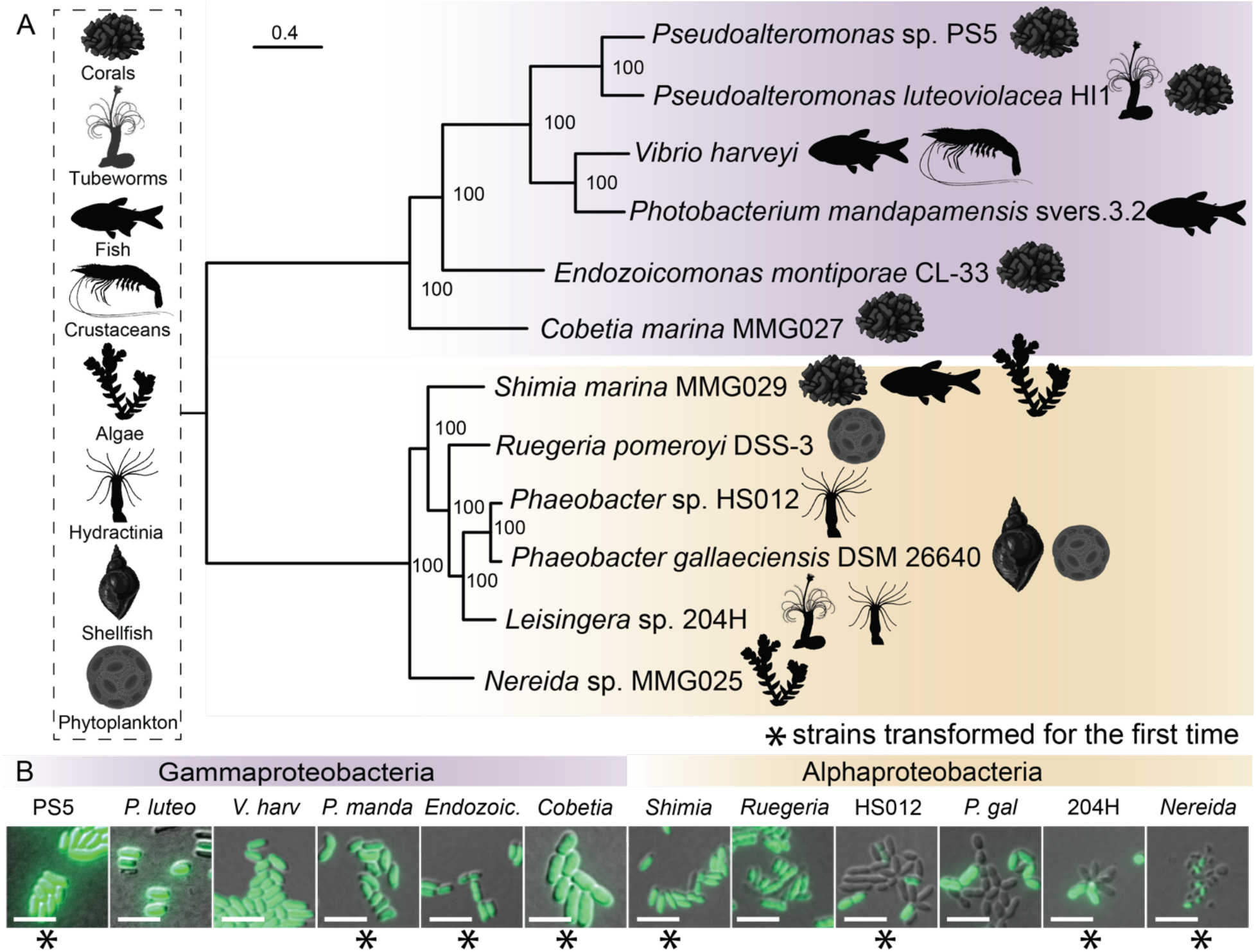
Diverse marine Proteobacteria are amenable to plasmid uptake and stable replication of toolkit plasmids. **(A)** Maximum likelihood whole genome phylogeny of 12 strains selected for manipulation and successfully transformed in this study (52, 53). All strains used in this study are known for their interaction with a range of marine biota and the icons depicting their associated host are shown in the vertical box. Gammaproteobacteria strains are highlighted in purple and Alphaproteobacteria strains are shown in gold. Scale bar is 0.4 and bootstraps were generated using the rapid-bootstrapping method (54). The tree was rooted at the midpoint with FigTree (v1.4.4) **(B)** Fluorescence microscopy of overnight cultures containing constitutively expressed RSF1010 *ori* fluorescence vector (CP25-*gfp*-T7). Scale bar is 5 µm.

## DISCUSSION

### Modular genetic tools provide insights about bacteria-stimulated metamorphosis

We tested a modular plasmid toolkit on a genetically tractable marine bacterium, *P. luteoviolacea*, that promotes the metamorphosis of the tubeworm *Hydroides elegans* (40, 41, 55) and produces several bioactive secondary metabolites (26, 29, 56, 57). We expand the tools available for functional interrogation of bacteria-stimulated metamorphosis in *P. luteoviolacea* by quantifying gene expression by a luminescence assay (Figure 1D and E), and using CRISPRi to knock down the secondary metabolite, violacein (Figure 2C and D), as well as a metamorphosis-associated gene, *macB* (Figure 3B) during the bacteria-tubeworm interaction. Distinct patterns of sheath (*macS*p) (41, 58) and baseplate (*macB*p) promoter induction suggest distinct mechanisms of gene regulation within the MACs gene cluster. Expression of the sheath gene was sensitive to bacterial mode of growth, while baseplate gene expression appeared static across the growth conditions tested. Although MACs are known to produce two effectors that stimulate tubeworm metamorphosis and kill eukaryotic cells (41, 58), the environmental conditions that promote MACs production remain poorly characterized. The tools developed here could help to characterize the conditions under which *P. luteoviolacea* MACs are produced or assembled and could help in the development of MACs or other contractile injection systems for use in biotechnology (59, 60). The modular tools in this work open new capabilities for interrogating bacterial biology, including the ability to quantify gene expression, knock down gene expression for rapid functional testing, and visualize bacteria during an *in vivo* interaction.

Whether, and how, bacteria and the animal are harmed or benefit from the interaction during bacteria-stimulated metamorphosis remains a prominent question in the field (38, 61, 62). Previous work by Gosselin *et al*. have shown that *Hydroides* is able to feed on bacteria as the sole food source (63). But until the present work, live bacteria within the gut of *Hydroides* juveniles had not been observed (Figure 3C) (21). The visualization of transgenic bacteria in *Hydroides* will enable future lines of research that can help dissect the role of microbiome seeding in bacteria-stimulated metamorphosis. More broadly, our results showcase the feasibility of using a modular plasmid toolkit to test hypotheses about bacteria-stimulated metamorphosis, and provides a framework for the interrogation of other bacteria and their products that promote host-microbe symbioses (36, 64, 65).

### Toolkit compatibility in diverse Proteobacteria and their potential for future study

In this work, we explore genetic tractability and gene function in 12 ecologically relevant marine Proteobacteria. These strains belong to two Proteobacterial classes, half of which were transformed for the first time (Figure 4). In the *Phaeobacter, Leisingera* and *Nereida* strains, expression of *gfp* was not uniformly observed in all cells imaged. However, this plasmid toolkit could be used in the future to identify promoters with that would express reporter genes in a greater proportion of the population. Compatibility with the broad-host-range plasmid backbone (RSF1010 origin of replication) suggests that all strains may be amenable to further manipulation with other toolkit coding sequences including luminescence reporters, complementation and CRISPRi.

The Gammaproteobacteria strains transformed in this study are a diverse selection of symbiosis-associated strains representing five genera (Figure 4A). To our knowledge, this is the first report of genetic tractability in strains from the genera *Endozoicomonas* and *Cobetia* (Figure 4B). *Endozoicomonas* species are among the most abundant bacterial symbionts in some corals and other marine hosts but are notoriously difficult to culture, therefore limiting our understanding of their functional roles in animal holobionts (66–68). Related strains of *Cobetia*, have been implicated in thermotolerance against bleaching in coral experiments with probiotic consortium treatments (69). The transformation of the representative *Endozoicomonas* and *Cobetia* strains in this study is a considerable step towards exploring function in coral host-microbiome interactions at a critical time to encourage the restoration of coral reefs (6, 70, 71). The genetic transformation of *Pseudoalteromonas* sp. PS5 in this study presents an opportunity to explore secondary metabolite production, including the coral metamorphosis-inducing compound, tetrabromopyrrole (Figure 4) (36). Other Gammaproteobacteria successfully transformed in this study include two bioluminescent strains, *Vibrio harveyi* and *Photobacterium mandapamensis* svers.3.2, which are associated with luminescence and organogenesis in squid (72, 73) and fish (74), respectively, but also as pathogens in aquaculture (75, 76) and corals (77, 78) (Figure 4A). In summary, the development of methods and established tractability of several new strains and genera have significant implications for the future of bacterial genetic development in established and emerging symbiosis systems.

The Alphaproteobacteria strains tested in this study fall within the *Roseobacter* group (Figure 4A), an ecologically important group of bacteria known to play a role in sulfur and carbon cycling on marine phytoplankton (79–81), in part due to their special capacity for lateral gene transfer and biofilm formation (82, 83). *Roseobacter* strains have also been explored as probiotics for the aquaculture industry (84–86). We explored the compatibility of the toolkit with the tractable, phytoplankton-associated species of *Phaeobacter gallaeciensis* (87), and *Ruegeria pomeroyi* (88), and demonstrate transformation for the first time with invertebrate microbiome-associated strains *Phaeobacter* sp. HS012 (89) and *Leisingera* sp. 204H (90) (Figure 4). The tractability of bacteria within the genus *Shimia* sp. has not previously been explored prior to this study, which may be of interest for coral microbiome studies in simulating impacted environments (91–94). Furthermore, there are no previous reports of the genetic manipulation of species in the *Nereida* genus, which have been isolated from kelp (95) and are associated with gall formations (96, 97). Tractability in this strain could help guide further understanding of microbe-seaweed interactions (98, 99), kelp aquaculture and the development of kelp probiotics (100). Taken together, the framework used in this study to establish genetic tractability can be used in future studies to explore the function of marine *Roseobacter* species in a wide range of symbiosis systems from the environment.

## CONCLUSION

The modular plasmid toolkit described here provides a basis for streamlining the genetic manipulation of marine bacteria for basic and applied purposes. These tools open up new possibilities to studying marine microbes in the context of plant and animal interactions, or with challenging and diverse non-model bacteria, ultimately helping us harness marine microbes for research, bioproduction and biotechnology.

## MATERIALS AND METHODS

### Bacterial Culture

A list of strains used in this study, isolation sources, accession numbers, and minimum inhibitory concentration can be found in Table S1. Environmental strains of marine bacteria were isolated and cultured on Marine Broth (MB) 2216 (BD Difco) and or natural seawater tryptone (NSWT) media (1 L 0.2 µm filtered natural seawater from Scripps Pier, La Jolla, CA, 2.5 g Tryptone, 1.5 g Yeast, 1.5 mL glycerol). MB and NSWT media are used interchangeably throughout the study; however, experiments were always conducted using only one media type. Marine bacteria were incubated between 25-30 °C, and cultures were shaken at 200 rpm. All liquid cultures were inoculated with a single colony and incubated between 16-18 hours, unless otherwise indicated. *E. coli* SM10*pir* and S17-1*pir* were cultured in LB (Miller, BD Difco) at 37 °C, shaking at 200 rpm. *E. coli* MFD*pir* (45) was cultured in LB supplemented with 0.3 mM Diaminopimelic acid (DAP). For *E. coli*, antibiotic selections with ampicillin, kanamycin, chloramphenicol were performed using a concentration of 100 µg/mL.

### Plasmid construction & Assembly

Golden Gate Assembly-compatible parts from the BTK, YTK (17, 18) and MMK used in this work can be found in Table S2. New plasmid parts were made by PCR amplifying insert and backbone fragments and combining them with Gibson Assembly with a 2:1 ratio (insert: backbone) (101). PCR amplification was performed with custom primers (Table S3), a high-fidelity DNA polymerase (PrimeSTAR GXL, Takara) and purified using a DNA Clean and Concentrator kit (Zymo Research). Part plasmids were assembled to make a stage 1 plasmid using Golden Gate Assembly, with T4 DNA ligase (Promega) and either BsaI or BsmBI (New England Biolabs), depending on the construct. Single-tube assembly was performed by running the following thermocycler program (BsaI/BsmBI): 37/42 °C for 5 minutes, 16 °C for 5 minutes, repeat 30x, 37/55 °C for 10 minutes, 80 °C for 10 minutes. The assemblies were directly electroporated into S17-1*pir* cells, confirmed by colony PCR (EconoTaq PLUS Green, LGC Biosearch) with internal primers and then shuttled to MFD*pir* cells for conjugation. The Ptac-sgRNA part plasmid with guide RNA was created to ensure expression of the sgRNA in *P. luteoviolacea*. To increase plasmid assembly efficacy, a BsmBI recognition site was moved to include the *bla* ampicillin resistance gene within the dCas9 part, enabling dual selection for positively assembled clones with kanamycin and ampicillin resistance. The CRISPRi assemblies were electroporated directly into SM10*pir* cells and shuttled to MFD*pir* cells for conjugation.

### Biparental conjugation in marine bacteria

*E. coli* donor strains (MFD*pir* or SM10*pir*) containing the mobilizable plasmids were grown under antibiotic selection in LB with the appropriate supplements (including 0.3 mM DAP for *E. coli* MFD*pir*). Conjugations were performed as previously described (17) with modifications for culturing marine bacteria. Briefly, several colonies of the recipient strains were inoculated and grown overnight in liquid culture. Recipient and donor cultures were spun down (4000 x g for 2 minutes) in a 1:1 OD_600_ ratio. All donor supernatant was removed leaving only the cell pellet. All but 100 µL of the recipient supernatant is removed and the cell pellet is resuspended. The recipient suspension is transferred to the donor pellet, which is resuspended with the recipient cells. Two 50 µL spots are plated onto NSWT (supplemented with 0.3 mM DAP for MFD-mediated conjugations). Spots are resuspended in 500 µL of liquid marine media and 100 µL is plated onto marine media containing antibiotic selection, according to the minimum inhibitory concentration (Table S1) Several of the bacteria take longer to grow or do not reach a high optical density (i.e. *Endozoicomonas, Ruegeria, Nereida*) in culture. Slower-growing marine bacteria were conjugated by growing larger 50 mL initial volumes of culture and spinning down the entire culture to reach 1:1 donor: host ratios.

### Phylogeny

Strains or close representative strains used in this study were compiled into a genome group on PATRIC v3.6.12 (102). A whole genome phylogenetic codon tree composed of 100 single copy genes (103) was performed using the Phylogenetic Tree Service (104–106). A Maximum likelihood phylogeny was generated using the best protein model found by RaxMLv8.2.11 (107), which was LG. Bootstraps were generated using the rapid bootstrapping algorithm with the default of 100 resamples (54). The tree was visualized with FigTree v1.4.4. and was rooted at the mid-line.

### Luciferase Culture and Assay

*P. luteoviolacea* containing plasmids with constitutive or native promoters driving *Nanoluciferase* (*Nluc*) were inoculated into 5 mL of MB or NSWT media with appropriate antibiotics and grown at 25 °C at 200 rpm for 24 hours. Each biological replicate was represented by a separate culture. Cultures used for the growth phase assay were inoculated as a 1:100 dilution with the appropriate antibiotic, and then incubated at 25 °C and shaking at 200 rpm. The luminescence of cultures was measured at exponential (OD_600_ of 0.35-1.0), early stationary (OD_600_ 1.0-1.45) or late stationary (OD_600_ 2.38-2.54) phases. For biofilm cultures, 1.5 mL of stationary-phase culture was pelleted and plated as a single spot on NSWT or MB plates. Biofilm plates were incubated at 20-25 °C for 24-28 hours. Each spot was scraped with a pipette tip and resuspended in 200 µL of NSWT or MB media before being resuspended in NSWT or MB. Luciferase reactions were performed with 100 µL of bacterial culture or biofilm resuspension aliquoted into opaque white walled 96-well plates (Corning #3642), with a modified protocol as written for Promega Nano-Glo Live Cell Assay System (Promega cat#N2011). Briefly, bacteria and the final reagent mix were read at a 1:1 ratio. Luminescence was measured on a Molecular Devices Microplate FilterMax F5 reader with a custom program on the Softmax Pro 7 software. Readings were done on the kinetic luminescence mode at 2-minute intervals for 20 minutes in total, using a 400 ms integration time, a 1 mm height read, and no other optimization or shaking settings. The detection limit is defined as three standard deviations above nine biological and technical replicates of WT *P. luteoviolacea*. Raw data were normalized to the OD_600_ of the culture used and plotted with an N=3 biological replicates.

### Violacein extraction

The specified *P. luteoviolacea* strains were struck onto NSWT media containing 200 µg/mL of streptomycin and kanamycin and incubated overnight at 25 °C. Single colonies were inoculated into 5 mL of liquid media containing the same antibiotic concentrations. Cultures were incubated at 25 °C, shaking at 200 rpm between 18 and 20 hours. Cultures were removed from the incubator and standardized to an OD_600_ of 1.5. The cells were pelleted and the supernatant was removed. The cell pellet was resuspended in 200 µL of 100% ethanol. The resuspended cells were pelleted and the supernatant containing the crude extract was recorded on a Biotek Synergy HT plate reader (Vermont, USA) using the Gen5 program (v2.00.18) with an endpoint reading at 580 nm.

### Microscopy

Microscopy was performed using a Zeiss Axio Observer.Z1 inverted microscope equipped with an Axiocam 506 mono camera and Neofluar10x/0.3 Ph1/DICI (*Hydroides* co-cultures) or Apochromat 100x/1.4 Oil DICIII (bacteria only) objectives. The Zeiss HE eGFP filter set 38 was used to capture GFPoptim-1 expression and Zeiss HE mRFP filter set 63 was used to capture *mRuby2* expression. For *Nanoluciferase* controls, images were captured using the same fluorescence exposure times as the *gfp* optim-1 and *mRuby2* labeled strains of the same species. Bacterial culture (2 μL) were added to freshly prepared 1% saltwater low-melt agarose (Apex catalog #20-103, Bioresearch products) pads on glass slides and coverslips were placed on top. *Hydroides elegans* were prepared in visualization chambers (Lab-Tek Chambered Coverglasses catalog #155411PK) with bacteria and imaged.

### Hydroides elegans culture

*Hydroides elegans* adults were collected from Quivira Basin, San Diego, California. The larvae were cultured and reared as previously described (40, 108). Larvae were maintained in beakers containing filtered artificial seawater (35 PSU) and were given new beakers with water changes daily. The larvae were fed live *Isochrysis galbana* and cultures were maintained as described previously. The larvae were used for metamorphosis assays once they reached competency (between 5 and 7 days old) (109).

### Hydroides elegans metamorphosis assays

Biofilm metamorphosis assays were performed using previously described methods (39, 40, 110). Briefly, bacteria were struck onto Marine Broth plates with 300 µg/mL kanamycin as appropriate and were incubated overnight at 25 °C. Up to 3 single colonies were inoculated into liquid broth and incubated overnight (between 15 and 18 hours), shaking at 200 rpm. Cultures were pelleted at 4000 g for 2 minutes, the spent media was removed and the cell pellets were washed twice with filtered artificial sea water (ASW). The concentration of the cells was diluted to OD_600_ of 0.1 and four 100 µL aliquots of the cell concentrate were added to 96-well plates. The cells were given between 2 and 3 hours to form biofilms, then the planktonic cells were removed and the adhered cells were washed twice with filtered ASW. Between 20 and 40 larvae were added to each well in 100 µL of filtered ASW. Metamorphosis was scored after 24 hours. Three biological replicates were performed on different days using separate *Hydroides* larvae originating from different male and female animals.

Chambered metamorphosis assays were performed using the same preparation principles as described above with the following modifications. Visualization chambers (Lab-Tek, Cat# 155411) were used for setting up the metamorphosis assay, then subsequently imaged. Inductive strains containing constitutively expressed *gfp*/*mRuby*/*Nluc* plasmids were struck out onto MB media containing 300 µg/mL kanamycin. Several colonies were inoculated into 5 mL MB media with antibiotics. Cultures were grown for 18 hours and cells were washed and allowed to form biofilms as described above. Cell concentrations ranging between OD_600_ 0.1 and 0.5 were used to elicit optimal metamorphosis. Larvae were concentrated and the resident filtered ASW was treated with 300 µg/mL kanamycin. Larvae were imaged 24 hours later.

### Online protocols

Selected protocols used in this study can be accessed on the Shikuma Lab protocols.io page: https://www.protocols.io/workspaces/shikuma-lab-sdsu

## ACKNOWLEDGEMENTS

We thank current and former Shikuma Lab members that helped with cloning and providing feedback for this paper including Taylor Darby, Milagros Esmerode, Nicole Jacobson, and Dr. Kate Nesbit. We thank Dr. Alison Gould, Dr. Stanley Maloy and Dr. Kristen Marhaver for their contribution of strains or samples for this study. We thank Dr. Alyssa Demko, Dr. Jennifer Sneed, Dr. Jennifer Doudna, Dr. Valerie Paul and Dr. Benjamin Rubin for their feedback on the project and manuscript. Schematic figures were created in part using Biorender.com.

This work was supported by the National Science Foundation (2017232404, A.T.A.; 1942251, N.J.S.; OCE-1837116, B.S.M.), the Gordon and Betty Moore Foundation (GBMF9344 to N.J.S.; https://doi.org/10.37807/GBMF9344), Office of Naval Research (N00014-20-1-2120 to N.J.S.), the National Institutes of Health, NIGMS (R35GM146722 to N.J.S.; R01ES030316 to B.S.M.) and the Alfred P. Sloan Foundation, Sloan Research Fellowship (N.J.S.). A.T.A. and N.J.S. are coinventors on provisional U.S. patent application Serial number 63/323,653, entitled “Genetic Engineering of Marine Bacteria for Biomaterial Production, Probiotic Use in Aquaculture and Marine Environmental Restoration” and assigned to San Diego State University Research Foundation.

---

**Table S1.**
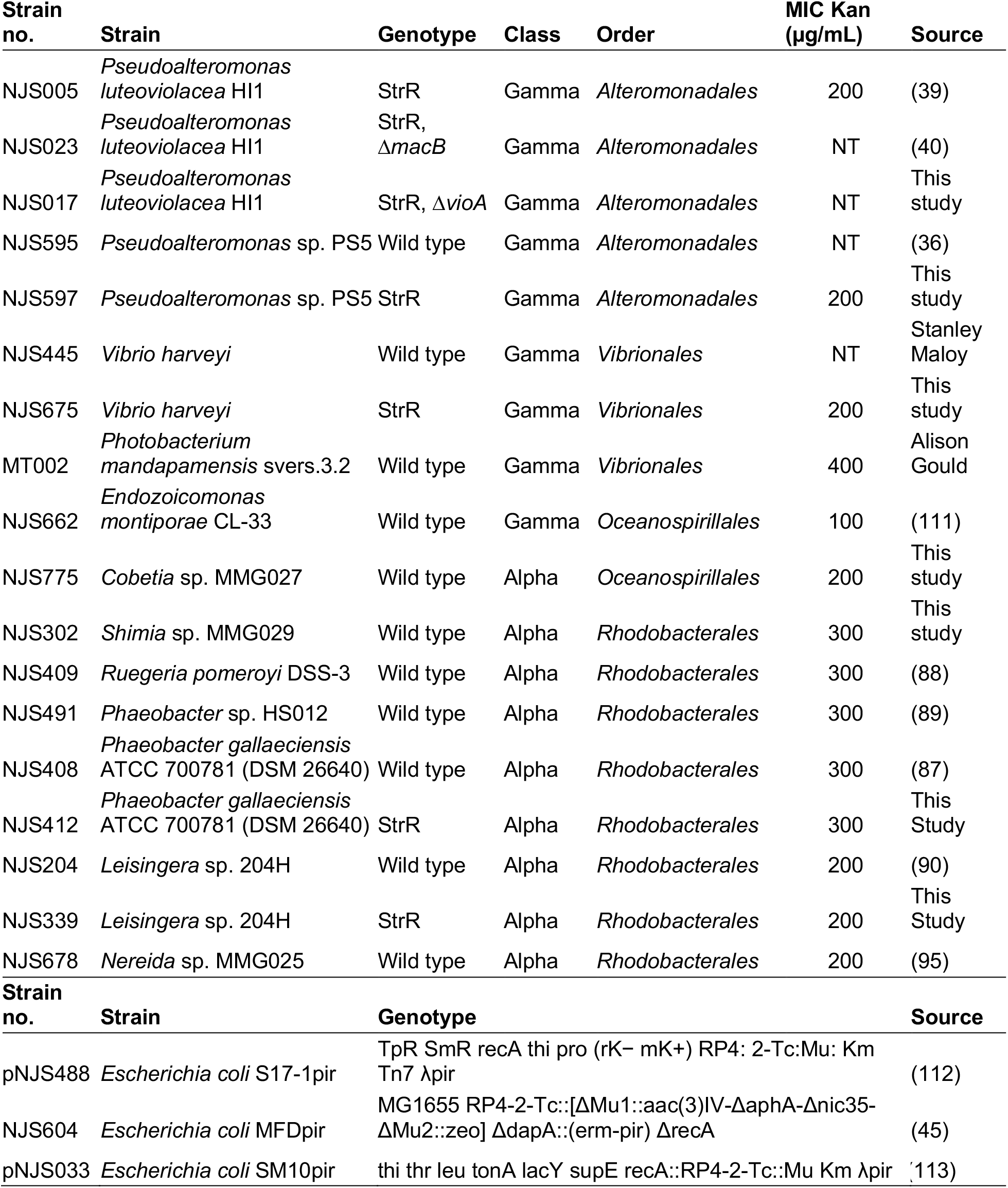
List of strains used in this study. MIC = minimum inhibitory concentration. Kan = kanamycin. Str = streptomycin. NT = antibiotic sensitivity not tested.

**Table S2.**
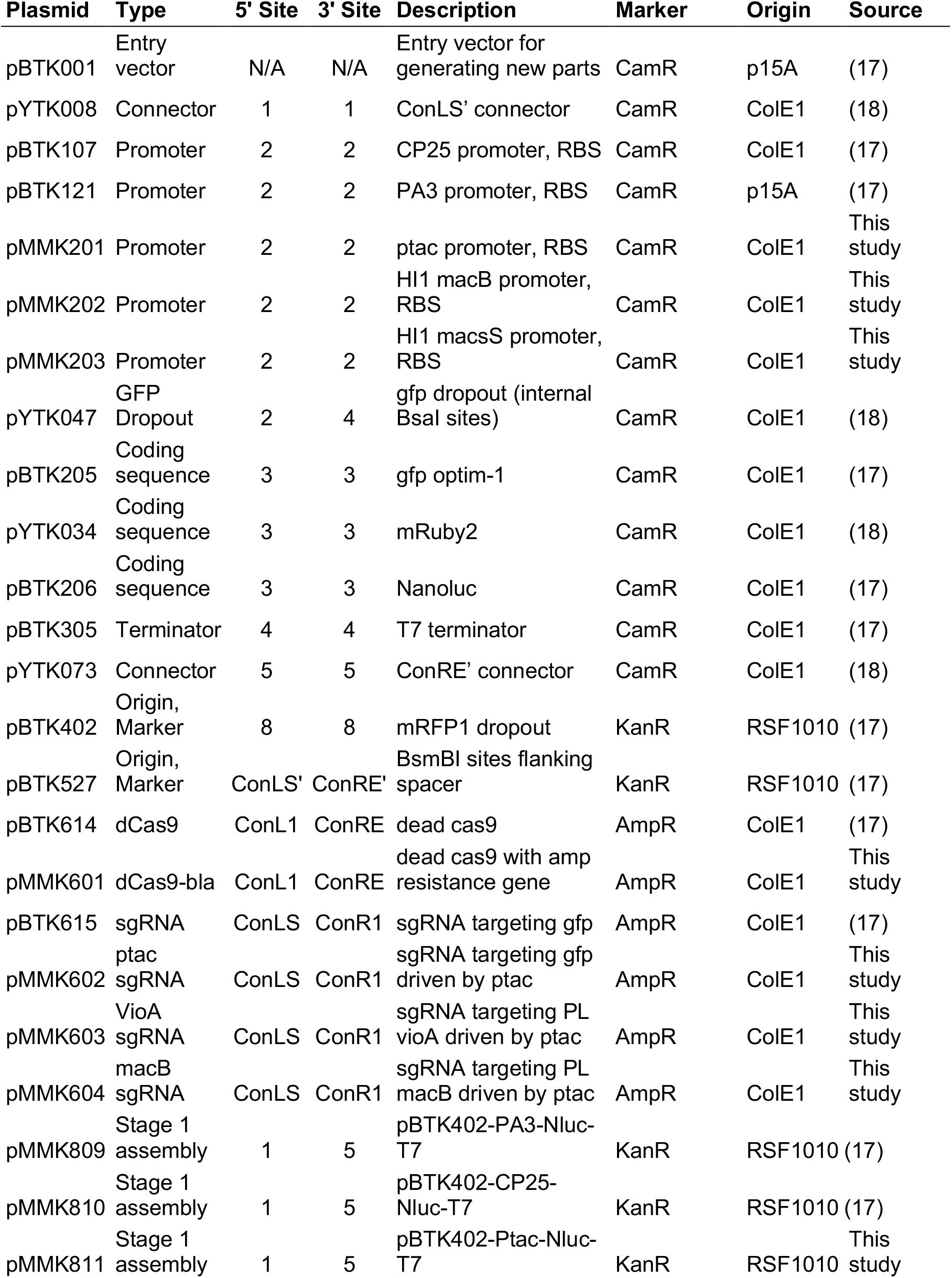
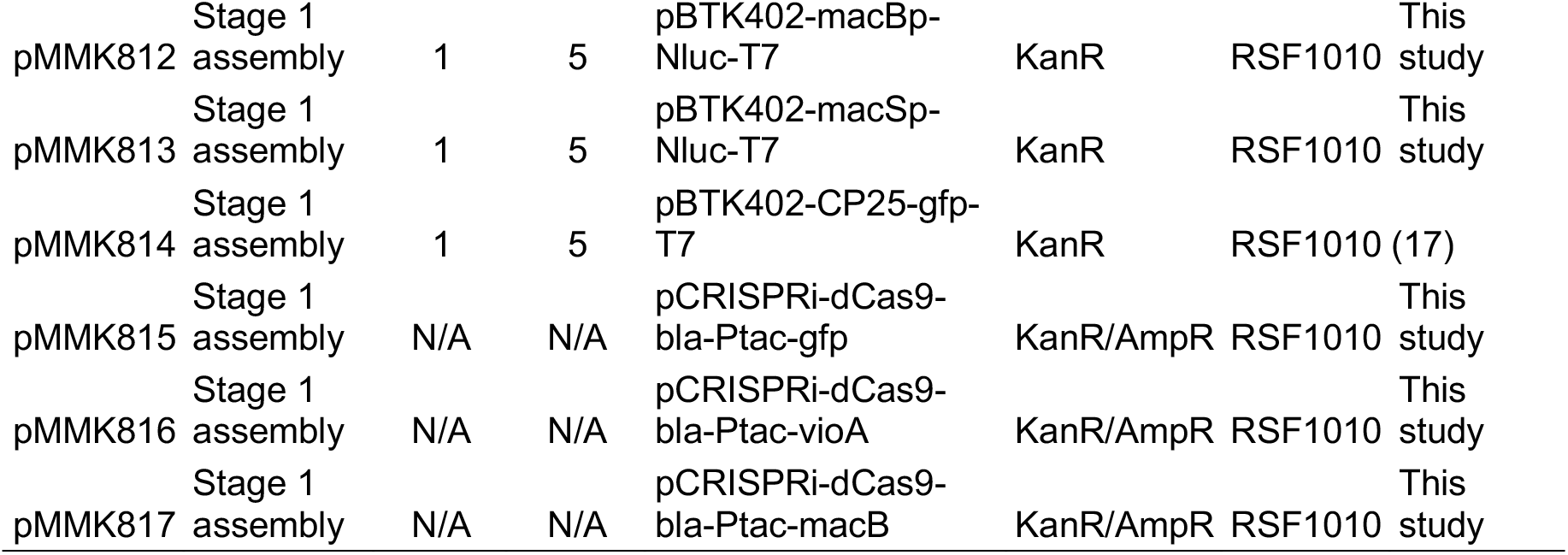
List of plasmids used in this study. N/A = 5’ or 3’ restriction site not applicable. Amp = ampicillin. Kan = kanamycin. Str = streptomycin.

**Table S3.**
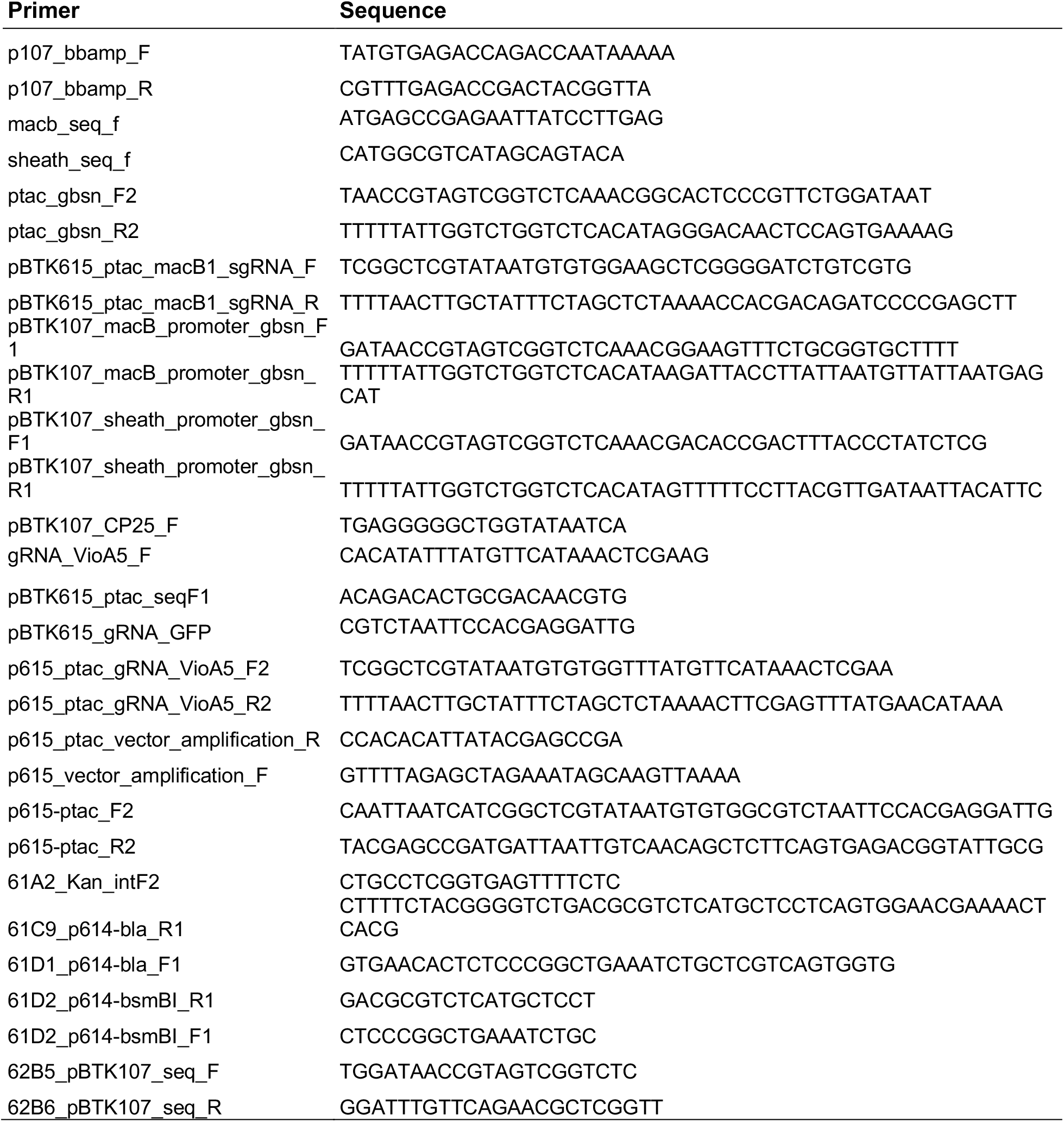
List of Primers used in this study.

## REFERENCES

1. Prado S, Romalde JL, Barja JL. 2010. Review of probiotics for use in bivalve hatcheries. Vet Microbiol 145:187–197.

2. D’Alvise PW, Lillebø S, Prol-Garcia MJ, Wergeland HI, Nielsen KF, Bergh Ø, Gram L. 2012. Phaeobacter gallaeciensis reduces vibrio anguillarum in cultures of microalgae and rotifers, and prevents vibriosis in cod larvae. PLoS One 7:e43996.

3. Peixoto RS, Rosado PM, Leite DC de A, Rosado AS, Bourne DG. 2017. Beneficial microorganisms for corals (BMC): Proposed mechanisms for coral health and resilience. Front Microbiol 8:341.

4. Kracke F, Vassilev I, Krömer JO. 2015. Microbial electron transport and energy conservation - The foundation for optimizing bioelectrochemical systems. Front Microbiol 6:575.

5. Nozzi NE, Oliver JWK, Atsumi S. 2013. Cyanobacteria as a Platform for Biofuel Production. Front Bioeng Biotechnol 0:7.

6. Peixoto RS, Voolstra CR, Sweet M, Duarte CM, Carvalho S, Villela H, Lunshof JE, Gram L, Woodhams DC, Walter J, Roik A, Hentschel U, Thurber RV, Daisley B, Ushijima B, Daffonchio D, Costa R, Keller-Costa T, Bowman JS, Rosado AS, Reid G, Mason CE, Walke JB, Thomas T, Berg G. 2022. Harnessing the microbiome to prevent global biodiversity loss. Nat Microbiol 1–10.

7. Lozada M, Dionisi HM. 2015. Microbial Bioprospecting in Marine Environments. Springer Handb Mar Biotechnol 307–326.

8. Paulsen SS, Strube ML, Bech PK, Gram L, Sonnenschein EC. 2019. Marine Chitinolytic Pseudoalteromonas Represents an Untapped Reservoir of Bioactive Potential. mSystems 4.

9. Madsen EL. 2011. Microorganisms and their roles in fundamental biogeochemical cycles. Curr Opin Biotechnol 22:456–464.

10. McFall-Ngai M, Hadfield MG, Bosch TCG, Carey H V, Domazet-Lošo T, Douglas AE, Dubilier N, Eberl G, Fukami T, Gilbert SF, Hentschel U, King N, Kjelleberg S, Knoll AH, Kremer N, Mazmanian SK, Metcalf JL, Nealson K, Pierce NE, Rawls JF, Reid A, Ruby EG, Rumpho M, Sanders JG, Tautz D, Wernegreen JJ. 2013. Animals in a bacterial world, a new imperative for the life sciences. Proc Natl Acad Sci U S A.

11. Paoli L, Ruscheweyh H-J, Forneris CC, Hubrich F, Kautsar S, Bhushan A, Lotti A, Clayssen Q, Salazar G, Milanese A, Carlström CI, Papadopoulou C, Gehrig D, Karasikov M, Mustafa H, Larralde M, Carroll LM, Sánchez P, Zayed AA, Cronin DR, Acinas SG, Bork P, Bowler C, Delmont TO, Gasol JM, Gossert AD, Kahles A, Sullivan MB, Wincker P, Zeller G, Robinson SL, Piel J, Sunagawa S. 2022. Biosynthetic potential of the global ocean microbiome. Nat 2022 6077917 607:111–118.

12. Sunagawa S, Coelho LP, Chaffron S, Kultima JR, Labadie K, Salazar G, Djahanschiri B, Zeller G, Mende DR, Alberti A, Cornejo-Castillo FM, Costea PI, Cruaud C, D’Ovidio F, Engelen S, Ferrera I, Gasol JM, Guidi L, Hildebrand F, Kokoszka F, Lepoivre C, Lima-Mendez G, Poulain J, Poulos BT, Royo-Llonch M, Sarmento H, Vieira-Silva S, Dimier C, Picheral M, Searson S, Kandels-Lewis S, Boss E, Follows M, Karp-Boss L, Krzic U, Reynaud EG, Sardet C, Sieracki M, Velayoudon D, Bowler C, De Vargas C, Gorsky G, Grimsley N, Hingamp P, Iudicone D, Jaillon O, Not F, Ogata H, Pesant S, Speich S, Stemmann L, Sullivan MB, Weissenbach J, Wincker P, Karsenti E, Raes J, Acinas SG, Bork P. 2015. Structure and function of the global ocean microbiome. Science (80-) 348.

13. Shetty RP, Endy D, Knight TF. 2008. Engineering BioBrick vectors from BioBrick parts. J Biol Eng 2:1–12.

14. Engler C, Kandzia R, Marillonnet S. 2008. A One Pot, One Step, Precision Cloning Method with High Throughput Capability. PLoS One 3:e3647.

15. Wiles TJ, Wall ES, Schlomann BH, Hay EA, Parthasarathy R, Guillemin K. 2018. Modernized tools for streamlined genetic manipulation and comparative study of wild and diverse proteobacterial lineages. MBio 9.

16. Vasudevan R, Gale GAR, Schiavon AA, Puzorjov A, Malin J, Gillespie MD, Vavitsas K, Zulkower V, Wang B, Howe CJ, Lea-Smith DJ, McCormick AJ. 2019. CyanoGate: A Modular Cloning Suite for Engineering Cyanobacteria Based on the Plant MoClo Syntax. Plant Physiol 180:39–55.

17. Leonard SP, Perutka J, Powell JE, Geng P, Richhart DD, Byrom M, Kar S, Davies BW, Ellington AD, Moran NA, Barrick JE. 2018. Genetic Engineering of Bee Gut Microbiome Bacteria with a Toolkit for Modular Assembly of Broad-Host-Range Plasmids. ACS Synth Biol 7:1279–1290.

18. Lee ME, DeLoache WC, Cervantes B, Dueber JE. 2015. A Highly Characterized Yeast Toolkit for Modular, Multipart Assembly. ACS Synth Biol 4:975–986.

19. Whitaker WR, Shepherd ES, Sonnenburg JL, Trehan I, Dominguez-Bello MG, Contreras M, Magris M, Hidalgo G, Baldassano RN, Anokhin AP, et al. 2017. Tunable Expression Tools Enable Single-Cell Strain Distinction in the Gut Microbiome. Cell 169:538--546.e12.

20. Buijs Y;, Bech PK, Vazquez Albacete D, Bentzon-Tilia M;, Sonnenschein EC, Gram LC;, Zhang S-D; 2022. Marine Proteobacteria as a source of natural products: advances in molecular tools and strategies. Nat Prod Rep 36:1333–1350.

21. Vijayan N, Lema KA, Nedved BT, Hadfield MG. 2019. Microbiomes of the polychaete Hydroides elegans (Polychaeta: Serpulidae) across its life-history stages. Mar Biol 166:19.

22. Bourne DG, Dennis PG, Uthicke S, Soo RM, Tyson GW, Webster N. 2013. Coral reef invertebrate microbiomes correlate with the presence of photosymbionts. ISME J 2013 77 7:1452–1458.

23. Stephens WZ, Burns AR, Stagaman K, Wong S, Rawls JF, Guillemin K, Bohannan BJM. 2015. The composition of the zebrafish intestinal microbial community varies across development. ISME J 2016 103 10:644–654.

24. Bourne DG, Morrow KM, Webster NS. 2016. Insights into the Coral Microbiome: Underpinning the Health and Resilience of Reef Ecosystems. Annu Rev Microbiol 70:317–340.

25. Bowman JP. 2007. Bioactive compound synthetic capacity and ecological significance of marine bacterial genus Pseudoalteromonas. Mar Drugs 5:220–241.

26. Maansson M, Vynne NG, Klitgaard A, Nybo JL, Melchiorsen J, Nguyen DD, Sanchez LM, Ziemert N, Dorrestein PC, Andersen MR, Gram L. 2016. An Integrated Metabolomic and Genomic Mining Workflow To Uncover the Biosynthetic Potential of Bacteria. mSystems 1:e00028–15.

27. Offret C, Desriac F, Le Chevalier P, Mounier J, Jégou C, Fleury Y. 2016. Spotlight on antimicrobial metabolites from the marine bacteria Pseudoalteromonas: Chemodiversity and ecological significance. Mar Drugs.

28. Chau R, Pearson LA, Cain J, Kalaitzis JA, Neilan BA. 2021. A Pseudoalteromonas Clade with Remarkable Biosynthetic Potential. Appl Environ Microbiol 87:1–16.

29. Thøgersen MS, Delpin MW, Melchiorsen J, Kilstrup M, Månsson M, Bunk B, Spröer C, Overmann J, Nielsen KF, Gram L. 2016. Production of the bioactive compounds violacein and indolmycin is conditional in a maeA mutant of Pseudoalteromonas luteoviolacea S4054 lacking the malic enzyme. Front Microbiol 7:1–11.

30. Carpizo-Ituarte E, Hadfield MG. 1998. Stimulation of metamorphosis in the polychaete Hydroides elegans Haswell (Serpulidae). Biol Bull 194:14–24.

31. Huang S, Hadfield MG. 2003. Composition and density of bacterial biofilms determine larval settlement of the polychaete Hydroides elegans. Mar Ecol Prog Ser 260:161–172.

32. Unabia CRC, Hadfield MG. 1999. Role of bacteria in larval settlement and metamorphosis of the polychaete Hydroides elegans. Mar Biol 133:55–64.

33. Tran C, Hadfield MG. 2011. Larvae of Pocillopora damicornis (Anthozoa) settle and metamorphose in response to surface-biofilm bacteria. Mar Ecol Prog Ser 433:85–96.

34. Negri AP, Webster NS, Hill RT, Heyward AJ. 2001. Metamorphosis of broadcast spawning corals in response to bacteria isolated from crustose algae. Mar Ecol Prog Ser 223:121–131.

35. Tebben J, Tapiolas DM, Motti CA, Abrego D, Negri AP, Blackall LL, Steinberg PD, Harder T. 2011. Induction of larval metamorphosis of the coral Acropora millepora by tetrabromopyrrole isolated from a Pseudoalteromonas bacterium. PLoS One 6:e19082.

36. Sneed JM, Sharp KH, Ritchie KB, Paul VJ. 2014. The chemical cue tetrabromopyrrole from a biofilm bacterium induces settlement of multiple Caribbean corals. Proc R Soc B Biol Sci 281.

37. Cavalcanti GS, Alker AT, Delherbe N, Malter KE, Shikuma NJ. 2020. The Influence of Bacteria on Animal Metamorphosis. Annu Rev Microbiol 74:137–158.

38. Shikuma NJ. 2021. Bacteria-Stimulated Metamorphosis: an Ocean of Insights from Investigating a Transient Host-Microbe Interaction. mSystems 6:e00754.-21.

39. Huang Y, Callahan S, Hadfield MG. 2012. Recruitment in the sea: bacterial genes required for inducing larval settlement in a polychaete worm. Sci Rep 2.

40. Shikuma NJ, Pilhofer M, Weiss GL, Hadfield MG, Jensen GJ, Newman DK. 2014. Marine Tubeworm Metamorphosis Induced by Arrays of Bacterial Phage Tail-Like Structures. Science (80-) 343:529–533.

41. Ericson CF, Eisenstein F, Medeiros JM, Malter KE, Cavalcanti GS, Zeller RW, Newman DK, Pilhofer M, Shikuma NJ. 2019. A contractile injection system stimulates tubeworm metamorphosis by translocating a proteinaceous effector. Elife 8:1–19.

42. Malter KE, Esmerode M, Damba M, Alker AT, Forsberg EM, Shikuma NJ. 2022. Diacylglycerol, PKC and MAPK signaling initiate tubeworm metamorphosis in response to bacteria. Dev Biol 487:99–109.

43. Fürste JP, Pansegrau W, Frank R, Blöcker H, Scholz P, Bagdasarian M, Lanka E. 1986. Molecular cloning of the plasmid RP4 primase region in a multi-host-range tacP expression vector. Gene 48:119–131.

44. Meyer R. 2009. Replication and conjugative mobilization of broad host-range IncQ plasmids. Plasmid 62:57–70.

45. Ferrières L, Hémery G, Nham T, Guérout AM, Mazel D, Beloin C, Ghigo JM. 2010. Silent mischief: Bacteriophage Mu insertions contaminate products of Escherichia coli random mutagenesis performed using suicidal transposon delivery plasmids mobilized by broad-host-range RP4 conjugative machinery. J Bacteriol 192:6418–6427.

46. Siebenlist U. 1979. Nuckotide sequence of the three major early promoters of bacteriophage T7. Nucleic Acids Res 6:1895–1907.

47. Jensen PR, Hammer K. 1998. The sequence of spacers between the consensus sequences modulates the strength of prokaryotic promoters. Appl Environ Microbiol 64:82–87.

48. Lee AK, Falkow S. 1998. Constitutive and Inducible Green Fluorescent Protein Expression in Bartonella henselae. Infect Immun.

49. Qi LS, Larson MH, Gilbert LA, Doudna JA, Weissman JS, Arkin AP, Lim WA. 2013. Repurposing CRISPR as an RNA-guided platform for sequence-specific control of gene expression. Cell 152:1173–1183.

50. Larson MH, Gilbert LA, Wang X, Lim WA, Weissman JS, Qi LS. 2013. CRISPR interference (CRISPRi) for sequence-specific control of gene expression. Nat Protoc 8:2180–2196.

51. Balibar CJ, Walsh CT. 2006. In vitro biosynthesis of violacein from L-tryptophan by the enzymes VioA-E from Chromobacterium violaceum. Biochemistry 45:15444–15457.

52. Davis JJ, Wattam AR, Aziz RK, Brettin T, Butler R, Butler RM, Chlenski P, Conrad N, Dickerman A, Dietrich EM, Gabbard JL, Gerdes S, Guard A, Kenyon RW, MacHi D, Mao C, Murphy-Olson D, Nguyen M, Nordberg EK, Olsen GJ, Olson RD, Overbeek JC, Overbeek R, Parrello B, Pusch GD, Shukla M, Thomas C, Vanoeffelen M, Vonstein V, Warren AS, Xia F, Xie D, Yoo H, Stevens R. 2020. The PATRIC Bioinformatics Resource Center: Expanding data and analysis capabilities. Nucleic Acids Res 48:D606--D612.

53. Guindon S, Gascuel O. 2003. A Simple, Fast, and Accurate Algorithm to Estimate Large Phylogenies by Maximum Likelihood. Syst Biol 52:696–704.

54. Stamatakis A, Hoover P, Rougemont J. 2008. A Rapid Bootstrap Algorithm for the RAxML Web Servers. Syst Biol 57:758–771.

55. Shikuma NJ, Antoshechkin I, Medeiros JM, Pilhofer M, Newman DK, Medeiros JM, Pilhofer M, Newman DK. 2016. Stepwise metamorphosis of the tubeworm Hydroides elegans is mediated by a bacterial inducer and MAPK signaling. Proc Natl Acad Sci 113:10097–10102.

56. Busch J, Agarwal V, Schorn M, Machado H, Moore BS, Rouse GW, Gram L, Jensen PR. 2019. Diversity and distribution of the bmp gene cluster and its Polybrominated products in the genus Pseudoalteromonas. Environ Microbiol 21:1575–1585.

57. Agarwal V, El Gamal AA, Yamanaka K, Poth D, Kersten RD, Schorn M, Allen EE, Moore BS. 2014. Biosynthesis of polybrominated aromatic organic compounds by marine bacteria. Nat Chem Biol 10:640–647.

58. Rocchi I, Ericson CF, Malter KE, Zargar S, Eisenstein F, Pilhofer M, Beyhan S, Shikuma NJ. 2019. A Bacterial Phage Tail-like Structure Kills Eukaryotic Cells by Injecting a Nuclease Effector. Cell Rep 28:295--301.e4.

59. Jiang F, Shen J, Cheng J, Wang X, Yang J, Li N, Gao N, Jin Q. 2022. N-terminal signal peptides facilitate the engineering of PVC complex as a potent protein delivery system. Sci Adv 8:2343.

60. Xu J, Ericson CF, Lien YW, Rutaganira FUN, Eisenstein F, Feldmüller M, King N, Pilhofer M. 2022. Identification and structure of an extracellular contractile injection system from the marine bacterium Algoriphagus machipongonensis. Nat Microbiol 2022 73 7:397–410.

61. Freckelton M, Nedved BT. 2020. When does symbiosis begin? Bacterial cues necessary for metamorphosis in the marine polychaete Hydroides elegans, p. 1–15. In Cellular Dialogues in the Holobiont. CRC Press.

62. Aldred N, Nelson A. 2019. Microbiome acquisition during larval settlement of the barnacle Semibalanus balanoides. Biol Lett 15.

63. Gosselin LA, Qian PY. 1997. Can bacterivory alone sustain larval development in the polychaete Hydroides elegans and the barnacle Balanus amphitrite? Mar Ecol Prog Ser 161:93–101.

64. Freckelton ML, Nedved BT, Hadfield MG. 2017. Induction of Invertebrate Larval Settlement; Different Bacteria, Different Mechanisms? Sci Rep 7:42557.

65. Petersen LE, Kellermann MY, Nietzer S, Schupp PJ. 2021. Photosensitivity of the Bacterial Pigment Cycloprodigiosin Enables Settlement in Coral Larvae—Light as an Understudied Environmental Factor. Front Mar Sci 8.

66. Neave MJ, Apprill A, Ferrier-Pagès C, Voolstra CR. 2016. Diversity and function of prevalent symbiotic marine bacteria in the genus Endozoicomonas. Appl Microbiol Biotechnol 100.

67. Neave MJ, Michell CT, Apprill A, Voolstra CR. 2017. Endozoicomonas genomes reveal functional adaptation and plasticity in bacterial strains symbiotically associated with diverse marine hosts. Sci Rep 7:1–12.

68. Pogoreutz C, Rädecker N, Cárdenas A, Gärdes A, Wild C, Voolstra CR. 2018. Dominance of Endozoicomonas bacteria throughout coral bleaching and mortality suggests structural inflexibility of the Pocillopora verrucosa microbiome. Ecol Evol 8:2240–2252.

69. Rosado PM, Leite DCA, Duarte GAS, Chaloub RM, Jospin G, Nunes da Rocha U, P. Saraiva J, Dini-Andreote F, Eisen JA, Bourne DG, Peixoto RS. 2019. Marine probiotics: increasing coral resistance to bleaching through microbiome manipulation. ISME J 13:921.

70. Li J, Yang Q, Dong J, Sweet M, Zhang Y, Liu C, Zhang Y, Tang X, Zhang W, Zhang S. 2022. Microbiome Engineering: A Promising Approach to Improve Coral Health. Engineering https://doi.org/10.1016/j.eng.2022.07.010.

71. Damjanovic K, Blackall LL, Webster NS, Oppen MJH van. 2017. The contribution of microbial biotechnology to mitigating coral reef degradation. Microb Biotechnol 10:1236–1243.

72. Nyholm S V., McFall-Ngai MJ. 2021. A lasting symbiosis: how the Hawaiian bobtail squid finds and keeps its bioluminescent bacterial partner. Nat Rev Microbiol 10.

73. Visick KL, Stabb E V., Ruby EG. 2021. A lasting symbiosis: how Vibrio fischeri finds a squid partner and persists within its natural host. Nat Rev Microbiol 0123456789.

74. Gould AL, Dunlap P V. 2019. Shedding Light on Specificity: Population Genomic Structure of a Symbiosis Between a Coral Reef Fish and Luminous Bacterium. Front Microbiol 10:2670.

75. Zhang XH, He X, Austin B. 2020. Vibrio harveyi: a serious pathogen of fish and invertebrates in mariculture. Mar Life Sci Technol. Springer.

76. King RK, Flick Jr GJ, Pierson D, Smith SA, Boardman GD, Coale Jr CW. 2004. Identification of Bacterial Pathogens in Biofilms of Recirculating Aquaculture Systems. J Aquat Food Prod Technol 13:125–133.

77. Ushijima B, Videau P, Burger AH, Shore-Maggio A, Runyon CM, Sudek M, Aeby GS, Callahan SM. 2014. Vibrio coralliilyticus strain OCN008 is an etiological agent of acute montipora white syndrome. Appl Environ Microbiol 80:2102–2109.

78. Ushijima B, Richards GP, Watson MA, Schubiger CB, Häse CC. 2018. Factors affecting infection of corals and larval oysters by Vibrio coralliilyticus. PLoS One 13:e0199475.

79. Dittmann KK, Sonnenschein EC, Egan S, Gram L, Bentzon-Tilia M. 2019. Impact of Phaeobacter inhibens on marine eukaryote-associated microbial communities. Environ Microbiol Rep 11:401–413.

80. Bramucci AR, Labeeuw L, Orata FD, Ryan EM, Malmstrom RR, Case RJ. 2018. The bacterial symbiont Phaeobacter inhibens Shapes the life history of its algal host emiliania huxleyi. Front Mar Sci 5:188.

81. Majzoub ME, Beyersmann PG, Simon M, Thomas T, Brinkhoff T, Egan S. 2019. Phaeobacter inhibens controls bacterial community assembly on a marine diatom. FEMS Microbiol Ecol 95:60.

82. Luo H, Moran MA. 2014. Evolutionary Ecology of the Marine Roseobacter Clade. Microbiol Mol Biol Rev 78:573–587.

83. Moran MA, Belas R, Schell MA, González JM, Sun F, Sun S, Binder BJ, Edmonds J, Ye W, Orcutt B, Howard EC, Meile C, Palefsky W, Goesmann A, Ren Q, Paulsen I, Ulrich LE, Thompson LS, Saunders E, Buchan A. 2007. Ecological genomics of marine Roseobacters. Appl Environ Microbiol 73:4559–4569.

84. Tesdorpf JE, Geers AU, Strube ML, Gram L, Bentzon-Tilia M. 2022. Roseobacter Group Probiotics Exhibit Differential Killing of Fish Pathogenic Tenacibaculum Species. Appl Environ Microbiol 88.

85. Sonnenschein EC, Jimenez G, Castex M, Gram L. 2021. The Roseobacter-Group Bacterium Phaeobacter as a Safe Probiotic Solution for Aquaculture. Appl Environ Microbiol 87:1–15.

86. Dittmann KK, Rasmussen BB, Melchiorsen J, Sonnenschein EC, Gram L, Bentzon-Tilia M. 2020. Changes in the microbiome of mariculture feed organisms after treatment with a potentially probiotic strain of Phaeobacter inhibens. Appl Environ Microbiol 86.

87. Ruiz-Ponte C, Cilia V, Lambert C, Nicolas JL. 1998. Roseobacter gallaeciensis sp. nov., a new marine bacterium isolated from rearings and collectors of the scallop Pecten maximus. Int J Syst Bacteriol 48 Pt 2:537–542.

88. González JM, Kiene RP, Moran MA. 1999. Transformation of sulfur compounds by an abundant lineage of marine bacteria in the alpha-subclass of the class Proteobacteria. Appl Environ Microbiol 65:3810–3819.

89. Deogaygay X, Delherbe N, Shikuma NJ. 2021. Draft Genome Sequences of Two Bacteria from the Roseobacter Group. Microbiol Resour Announc 10.

90. Cavalcanti GS, Wasserscheid J, Dewar K, Shikuma NJ. 2020. Complete Genome Sequences of Two Marine Biofilm Isolates, Leisingera sp. nov. Strains 201A and 204H, Novel Representatives of the Roseobacter Group. Microbiol Resour Announc 9.

91. Godwin S, Bent E, Borneman J, Pereg L. 2012. The Role of Coral-Associated Bacterial Communities in Australian Subtropical White Syndrome of Turbinaria mesenterina. PLoS One 7:e44243.

92. Apprill A, Marlow HQ, Martindale MQ, Rappé MS. 2009. The onset of microbial associations in the coral Pocillopora meandrina. ISME J 2009 36 3:685–699.

93. Zhang Y, Yang Q, Zhang Y, Ahmad M, Ling J, Tang X, Dong J. 2021. Shifts in abundance and network complexity of coral bacteria in response to elevated ammonium stress. Sci Total Environ 768:144631.

94. Silva DP, Villela HDM, Santos HF, Duarte GAS, Ribeiro JR, Ghizelini AM, Vilela CLS, Rosado PM, Fazolato CS, Santoro EP, Carmo FL, Ximenes DS, Soriano AU, Rachid CTCC, Vega Thurber RL, Peixoto RS. 2021. Multi-domain probiotic consortium as an alternative to chemical remediation of oil spills at coral reefs and adjacent sites. Microbiome 9:1–19.

95. Alker AT, Hern NA, Ali MA, Baez MI, Baswell BC, Baxter BI, Blitz A, Calimlim TM, Chevalier CA, Eguia CA, Esparza T, Fuller AE, Gwynn CJ, Hedin AL, Johnson RA, Kaur M, Laxina RT, Lee K, Maguire PN, Martelino IF, Melendez JA, Navarro JJ, Navarro JN, Osborn JM, Padilla MR, Peralta ND, Pureza JLR, Rojas JJ, Romo TR, Sakha M, Salcedo GJ, Sims KA, Trieu TH, Niesman IR, Shikuma NJ. 2022. Draft Genome Sequence of Nereida sp. Strain MMG025, Isolated from Giant Kelp. Microbiol Resour Announc 11:70–72.

96. Arahal DR, Pujalte MJ, Rodrigo-Torres L. 2016. Draft genomic sequence of Nereida ignava CECT 5292T, a marine bacterium of the family Rhodobacteraceae. Stand Genomic Sci 11:1–8.

97. Ashen JB, Goff LJ. 2000. Molecular and ecological evidence for species specificity and coevolution in a group of marine algal-bacterial symbioses. Appl Environ Microbiol 66:3024–3030.

98. Egan S, Harder T, Burke C, Steinberg P, Kjelleberg S, Thomas T. 2013. The seaweed holobiont: understanding seaweed–bacteria interactions. FEMS Microbiol Rev 37:462–476.

99. Singh RP, Reddy CRK. 2014. Seaweed-microbial interactions: Key functions of seaweed-associated bacteria. FEMS Microbiol Ecol 88:213–230.

100. Hofer U. 2021. A probiotic for seaweed. Nat Rev Microbiol 19:618.

101. Gibson DG, Young L, Chuang R-Y, Venter JC, Hutchison CA, Smith HO. 2009. Enzymatic assembly of DNA molecules up to several hundred kilobases. Nat Methods 6:343–345.

102. Wattam AR, Davis JJ, Assaf R, Boisvert S, Brettin T, Bun C, Conrad N, Dietrich EM, Disz T, Gabbard JL, Gerdes S, Henry CS, Kenyon RW, Machi D, Mao C, Nordberg EK, Olsen GJ, Murphy-Olson DE, Olson R, Overbeek R, Parrello B, Pusch GD, Shukla M, Vonstein V, Warren A, Xia F, Yoo H, Stevens RL. 2017. Improvements to PATRIC, the all-bacterial bioinformatics database and analysis resource center. Nucleic Acids Res 45:D535--D542.

103. Davis JJ, Gerdes S, Olsen GJ, Olson R, Pusch GD, Shukla M, Vonstein V, Wattam AR, Yoo H. 2016. PATtyFams: Protein Families for the Microbial Genomes in the PATRIC Database. Front Microbiol 7:118.

104. Edgar RC. 2004. MUSCLE: Multiple sequence alignment with high accuracy and high throughput. Nucleic Acids Res 32:1792–1797.

105. Cock PJA, Antao T, Chang JT, Chapman BA, Cox CJ, Dalke A, Friedberg I, Hamelryck T, Kauff F, Wilczynski B, De Hoon MJL. 2009. Biopython: freely available Python tools for computational molecular biology and bioinformatics. Bioinformatics 25:1422–1423.

106. Katoh K, Standley DM. 2013. MAFFT Multiple Sequence Alignment Software Version 7: Improvements in Performance and Usability. Mol Biol Evol 30:772–780.

107. Stamatakis A. 2014. RAxML version 8: a tool for phylogenetic analysis and post-analysis of large phylogenies. Bioinformatics 30:1312–1313.

108. Nedved BT, Hadfield MG. 2009. Hydroides elegans (Annelida: Polychaeta): A Model for Biofouling Research. Mar idustrial biofouling 4:203–217.

109. Nesbit KT, Shikuma N. Developmental staging of the complete life cycle of the model marine tubeworm Hydroides elegans https://doi.org/10.1101/2022.10.24.513551.

110. Alker AT, Delherbe N, Purdy TN, Moore BS, Shikuma NJ. 2020. Genetic examination of the marine bacterium Pseudoalteromonas luteoviolacea and effects of its metamorphosis-inducing factors. Environ Microbiol 22:4689–4701.

111. Yang C-S, Chen M-H, Arun AB, Chen CA, Wang J-T, Chen W-M. 2010. Endozoicomonas montiporae sp. nov., isolated from the encrusting pore coral Montipora aequituberculata. Int J Syst Evol Microbiol 60:1158–1162.

112. de Lorenzo V, Timmis KN. 1994. Analysis and construction of stable phenotypes in gram-negative bacteria with Tn5- and Tn10-derived minitransposons. Methods Enzymol 235:386–405.

113. Simon R, Priefer U, Pühler A. 1983. A Broad Host Range Mobilization System for In Vivo Genetic Engineering: Transposon Mutagenesis in Gram Negative Bacteria. Bio/Technology 1983 19 1:784–791.

